# Mapping the Spatial Proteome of Leukemia Cells Undergoing Fludarabine Treatment

**DOI:** 10.1101/2024.10.10.617576

**Authors:** Maria A. Globisch, Henrik Gezelius, Anna Pia Enblad, Anders Lundmark, Mariya Lysenkova Wiklander, Olga Krali, Martin Åberg, Arja Harila, Amanda Raine, Claes Andersson, Jessica Nordlund

## Abstract

Recent advancements in spatial biology have revolutionized our understanding of the organization and functional dynamics of cells and tissues. In this study, we applied Molecular Pixelation (MPX), a single-cell spatial proteomics assay, to investigate the modulation of the cell surface proteome in an in vitro drug screening model using the *ETV6::RUNX1* acute lymphoblastic leukemia (ALL) cell line, Reh. Specifically, we focused on the in vitro response to fludarabine, a chemotherapeutic agent used prior to allogenic stem cell transplantation and chimeric antigen receptor (CAR)-T cell therapy in high-risk, refractory, or relapsed ALL patients. Using MPX, we quantified changes in protein abundance, spatial distribution, and colocalization of 76 targeted cell surface proteins in Reh cells before and after fludarabine treatment. Our analysis revealed 25 proteins with altered abundance, 24 proteins with increased polarity, and 138 protein pairs with modified colocalization following treatment. Notably, the tetraspanins CD82 and CD53, which are known for their roles in chemotherapy resistance, exhibited increased abundance, polarization, and colocalization post-treatment, suggesting their potential as a therapeutic scaffold. These findings underscore the unique ability of spatially resolved single-cell proteomics to uncover nuanced cellular responses that would otherwise remain undetected.

## Introduction

Until recently, allogenic stem cell transplantation (allo-SCT) was the primary last-line treatment option for high-risk acute lymphoblastic leukemia (ALL) cases with insufficient response to first-line chemotherapy, treatment-refractory disease, or relapse^1,2^. However, the emergence of chimeric antigen receptor (CAR)-T cell therapy provides a more targeted and potent alternative, especially for refractory and relapsed cases. Despite its promise, the efficacy of CAR-T therapy remains variable, often hindered by challenges such as antigen evasion and CAR-T cell persistence^3,4^. While these obstacles are recognized^5^, the underlying mechanisms driving suboptimal CAR-T responses are still not fully understood, leaving critical gaps in our knowledge.

Fludarabine, a purine nucleotide analogue, is a critical component of preparative regimens for both CAR-T and allo-SCT therapies^6-8^. In this context, fludarabine enhances the efficacy of subsequent therapies through mechanisms such as immunosuppression, lymphodepletion, and tumor burden reduction, thereby creating more favorable microenvironment for these interventions^9,10^. By disrupting DNA synthesis and repair, fludarabine is well known to induce apoptosis in proliferating cancer cells^11^. Previous work has indicated a positive correlation between fludarabine exposure and improved outcomes in B-cell ALL patients receiving CAR-T therapy^12^. However, variable outcomes have been observed with intensified fludarabine exposure prior to allo-SCT ^13^. Thus, despite the essential role fludarabine plays as a preparatory treatment, our understanding of its impact on leukemia cells and the factors influencing its efficacy remain limited.

Spatial proteomics approaches enable the study of the spatial and temporal organization of the proteome in single cells, facilitating the identification of novel biomarkers and therapeutic targets^14^. As these methodologies evolve, they promise to advance our understanding of cancer biology and inform personalized treatment strategies. Previously, we coupled an in vitro drug screen with single-cell RNA-sequencing (scRNA-seq) to assess the transcriptional response of leukemia cells after fludarabine treatment^15^. We confirmed fludarabine’s effect on disrupting DNA synthesis and identified several differentially expressed genes coding for cell surface proteins ^15^, which suggested that this drug may modulate the cell-surface proteome.

Herein, we employ a single-cell spatial proteomics assay that quantifies protein abundance, spatial distribution, and colocalization of 76 targeted surface proteins using a DNA sequencing-based method^16^ to analyze the cell surface proteome of leukemia cells after fludarabine treatment. Through this approach, we identified a subset of significantly modulated proteins in leukemia cells post-treatment and demonstrate a proof-of-concept phenotypic response to fludarabine’s effect on leukemia cells.

## Materials and Methods

### Cell culture

The human *ETV6::RUNX1* Reh B-ALL cell-line (DSMZ #ACC 22;,) was maintained in RPMI-1640 medium (Sigma #R0883) supplemented with 10% heat-inactivated fetal bovine serum (HIFBS; Sigma #F9665), 2 mM L-Glutamine (Sigma #G7513), 100 U/mL Penicillin, and 100 µg/mL Streptomycin (Sigma #P0781). Cells were cultured at 37°C in a humidified chamber with 95% humidity and 5% CO_2_. The first two subcultures were supplemented with an additional HIFBS (20%) to enhance cell growth. Cells were split twice a week. The Reh cell line from DSMZ was verified by karyotyping^17^ and the Eurofins human STR profiling cell line authentication service^18^.

### Fludarabine treatment

The concentration of fludarabine was selected from the results of a previous study where the fluorometric microculture cytotoxicity assay (FMCA) was coupled with scRNA-seq to evaluate the transcriptional response to fludarabine in Reh cells^15^. In brief, the selected concentration of 0.56 µM fludarabine resulted in a survival index (SI) of approximately 50%, eliciting a cellular response, while maintaining sufficient surviving cells for downstream analysis^15^. Herein, Reh cells were treated with either: 0.56 µM fludarabine (Selleckchem #S1391) or dimethyl sulfoxide (DMSO, Honeywell #D5879) for 72 hours (hrs). DMSO served as a vehicle control as fludarabine was prepared in 100% DMSO prior to cell treatment and the final concentration of DMSO in all wells was 0.3%.

### Cell collection

After 72 hrs, cells growing in suspension were transferred to falcon tubes. The cell suspensions were spun at 200 RCF for 10 mins at room temperature (RT) and the supernatant was removed and cells were rinsed twice with Dulbecco’s Phosphate Buffered Saline (DPBS) and resuspended in 1 mL of DPBS. The concentration and viability of the cells was assessed with the AO DAPI Staining Solution 18 (ChemoMetec #910-3018) on a NucleoCounter-3000 (ChemoMetec) according to the manufacturer’s instructions. The viability of the cells at the time of collection was >90% (Supplementary Table 1).

### Single-cell spatial proteomics library preparation

One million cells from each condition were processed in duplicate for single-cell spatial proteomics (Molecular Pixelation, MPX) according to the user manual of the Single Cell Spatial Proteomics Kit, Immunology Panel I, Human (Pixelgen Technologies. #PXGIMM001). In summary, cells were fixed with 1% methanol-free paraformaldehyde (PFA, ThermoFisher #28906), rinsed with a wash buffer, blocked, frozen and stored at -80°C until further processing. The cells were thawed, washed, and incubated with 80 antibody oligonucleotide conjugates (AOCs). The 80 AOCs comprised of 76 cell surface proteins (Supplementary Table 2) and 4 control proteins (Supplementary Table 3). Cells with bound AOCs were then hybridized with single-stranded DNA molecules (DNA pixels A) that contained a concatemer of a UMI sequence called unique pixel identifier (UPI). DNA pixels A incorporated the UPI-A barcode onto the oligonucleotides of neighboring AOCs by a gap-fill ligation reaction, which created spatial protein neighborhoods on the cell surface. DNA pixels A were then degraded enzymatically (only the UPI-A barcode remained on the AOCs) and a second set of UPIs (UPI-B) were incorporated onto the AOCs with a second set of DNA pixels (DNA pixels B) in a similar manner to generate overlaps between spatial zones defined by UPI-A, which generate the spatial proteome map^16^. DNA pixels B were subsequently degraded enzymatically, leaving only the UPI-B barcodes on the AOCs. The samples were then indexed with unique primer sets (NEBNext Unique Dual Index Primers, New England Biolabs #E6441A) and the libraries were amplified by PCR. The AMPure XP Reagent (Beckman Coulter Life Sci. #A63882)) was used to purify the amplified PCR products with a 0.2 mL magnetic separator. The quality and concentrations of the libraries were evaluated with the High Sensitivity DNA ScreenTape Assay (Agilent Technologies #5067-5584/5) on a 2200 TapeStation) prior to sequencing. A summary of the number of cells fixed and used for each pixelation step per condition and replicate can be found in Supplementary Table 1.

### Single-cell spatial proteomic sequencing

The libraries were pooled to an equimolar concentration and sequenced with 15% PhiX spike-in on one lane of an S4 flowcell on a NovaSeq6000 Sequencer (Illumina) according to routine protocols at the SciLifeLab National Genomics Infrastructure in Uppsala, Sweden.

### Data analysis

#### MPX data processing and visualizations

The raw data (fastq files) were processed using the nf-core/pixelator pipeline (version 0.12.0)^19^ which filters out low-quality reads, maps the relative locations of AOCs to form >1,000 spatial zones per single-cell in 3D, and produces protein counts and spatial scores for downstream analyses. Downstream analyses were performed according to https://github.com/PixelgenTechnologies/pixelatorR. In short, low-quality cells were removed and protein counts were normalized (centered log ratio-transformed; CLR-transformed), the most variable markers were identified, and marker filtering was performed to remove markers that were not expressed by the assayed cells. Antibody markers with a lower median abundance than the isotype controls were removed. Dimensionality reduction (Uniform Manifold Approximation and Projection; UMAP) was used to visualize the normalized and scaled data. A two-sided Wilcoxon rank sum test was used to determine 1) differential abundance 2) differential polarity scores and 3) differential colocalization between the fludarabine treated and control (DMSO treated) Reh cells. For all analyses, a Benjamin-Hochberg (BH) adjusted (adj.) p-value < 0.05 was considered statistically significant and adj. p-values were categorized as following: not significant (ns) p>0.05, *p≤ 0.05, **p≤ 0.01, ***p≤ 0.001, and ****p≤ 0.0001.

#### Transcriptome data analysis and visualizations

The scRNA-seq data generated from fludarabine and DMSO treated Reh cells (10X Genomics Chromium Single Cell Gene Expression Flex; 10X-fixed) were obtained from GSE229617^15^. Single-cell transformed and normalized counts were used to visualize the data and a differential gene expression analysis was performed using a MAST test^20^. In the differential gene expression analysis, fludarabine treated cells were compared with DMSO control cells. The threshold for determining significant differentially expressed genes was log2FC > 0.50 and adj. p-value < 0.01. A scatter plot was made using the centered log ratio (CLR)-transformed data of the 25 differentially expressed proteins (MPX data) and the log normalized (aggregate) scRNA-seq data of genes coding for the corresponding cell surface proteins. The genes coding for the assayed cell surface proteins can be found on Supplementary Table 2. The correlation between the transcript counts (scRNA-seq) and protein abundance (MPX) was assessed using Pearson correlation.

### Immunocytochemistry (ICC)

Reh cells were cultured, seeded, treated, collected and counted as described above. One million cells were transferred to Eppendorf tubes and spun at 200 RCF for 10 min at RT. The supernatant was removed and the pellets were resuspended in 2% FBS diluted in PBS to a final concentration of 250,000 cells/mL. Slides were prepared for ICC by using a cytospin. In short, Superfrost Plus Slides (Thermo Fisher #J1800AMNZ) were attached to Epredia EZ Single Cytofunnels with White Filter Cards (Epredia, #A78710003). The cell solutions were added to cytofunnels and the samples were spun for 5 mins at 700 RPM in a Cytospin 4 centrifuge (Thermo Fisher). The slides were air-dried overnight at RT in a vertical position and the next day the cell smears were circled with a hydrophobic pen (Abcam #ab26-01) and fixed with 2% methanol-free PFA diluted in PBS or methanol (Fisher Scientific #A456-212) for 10 minutes at RT. Slides were air dried for 2 hrs at RT and then stored at -20°C until ready to process for ICC.

Slides were thawed for 5 mins at RT, rehydrated and permeabilized with 0.5% Triton-X diluted in PBS for 10 minutes at RT, and blocked with blocking buffer (PBS supplemented with 0.1% Triton-X, 2% BSA and 5% donkey serum) for 30 minutes at RT. Primary antibodies listed on Supplementary Table 4 were diluted in blocking buffer, added to the cell smears and incubated in a humidified chamber overnight at 4°C. Slides were then washed 3×5 mins with a washing buffer (PBS + 0.5% Tween-20). Secondary antibodies listed on Supplementary Table 5 were diluted in PBS and incubated for 1 hr at RT. Slides were then rinsed 3×5 mins with washing buffer and nuclei were stained with NucBlue Fixed Cell Stain Ready Probe Reagent (Invitrogen #R37606) was prepared following the manufacturer’s instructions and cells were incubated for 15 mins at RT. The slides were then mounted with Fluoromount-G (Invitrogen #00-4958-02). The slides were then left to dry for 48h at RT and then stored at 4°C until ready to image.

### Immunofluorescent imaging, analysis, and processing

Stained cells were imaged with a Zeiss LMS700 confocal microscope with 20X and 63X objectives at the BioVis Core Facility at Uppsala University. For comparison purposes, different sample images of the same antibodies were taken with the same acquisition settings. For image analysis, the open-source software ImageJ (https://imagej.net) was used. To measure the nuclei perimeter in Reh cells, a threshold was set on DAPI images (taken with a 20X objective) to distinguish nuclei from the background. The “analyze particle” function was then used to identify the number of nuclei in each image. The area and perimeter of all cells were calculated by Image J. The diameters of all nuclei were calculated by dividing the circular perimeter (circumference) by π. Python was used to conduct a statistical analysis (t-test) of nuclei diameter between fludarabine (n=124) and control (n=85) cells and then the data were visualized with boxplots. Only single cells were included in the analysis (cell clusters were excluded). To assemble representative immunofluorescent images, Adobe Photoshop was used to adjust the brightness and contrast of images, and for comparison purposes, the same adjustment was made for the same epitope of all samples.

## Results

### Study design and quality control analysis

To investigate the cell surface proteome of leukemia cells after exposure to fludarabine we utilized Molecular Pixelation (MPX), a spatial proteomics method that quantifies protein abundance, spatial distribution, and colocalization of targeted surface proteins^16^. The *ETV6::RUNX1* B-ALL cell line, Reh, was incubated with fludarabine or DMSO (vehicle control) for 72 hrs and processed for MPX in duplicate (Figure 1A). The nf-core/pixelator pipeline reported the number of identified cells in silico, average reads per cell, the average number of useable reads per cell, and the median antibody molecules per cell (Supplementary Table 6). Quality control analysis of the sequencing data was performed to filter out low quality cells (see methods). An edge-rank plot was used to visualize the range and distribution of antibody counts (edges) and the components (single cells) declined rapidly at around 20,000 edges, thus a cutoff was set at that point to filter out low-quality cells that deviated from the component size distribution (Figure 1B). After filtering, 1045-1178 cells (of a target 1000 cells), remained per replicate (Figure 1C). After normalization (CLR-transformation), 18 proteins with a lower median abundance than the MPX isotype controls (IgG1, IgG2a, and IgG2b) were removed (Supplementary Figure 1). The proteins below the control cut-off are proteins typically not expressed by cells of the B-cell lineage. The remaining proteins were used for downstream analyses. Of note, the fludarabine treated Reh cells had on average fewer mean reads, but more edges (UMIs) and A pixels (UPI-A), which resulted in fewer mean edges per A pixel (Figure 1D). As more UPI-A pixels would attach to larger cell surface areas, this suggested that the post-treatment Reh cells were larger in size, which has been speculated to occur in replicative and drug-induced senescence^21^. To further validate this observation, using ICC and Image J, we measured the diameter of nuclei as a proxy of cell size^22^ and confirmed that the fludarabine treated cells were indeed larger than DMSO treated control cells (Supplementary Figure 2).

**Figure 1.**
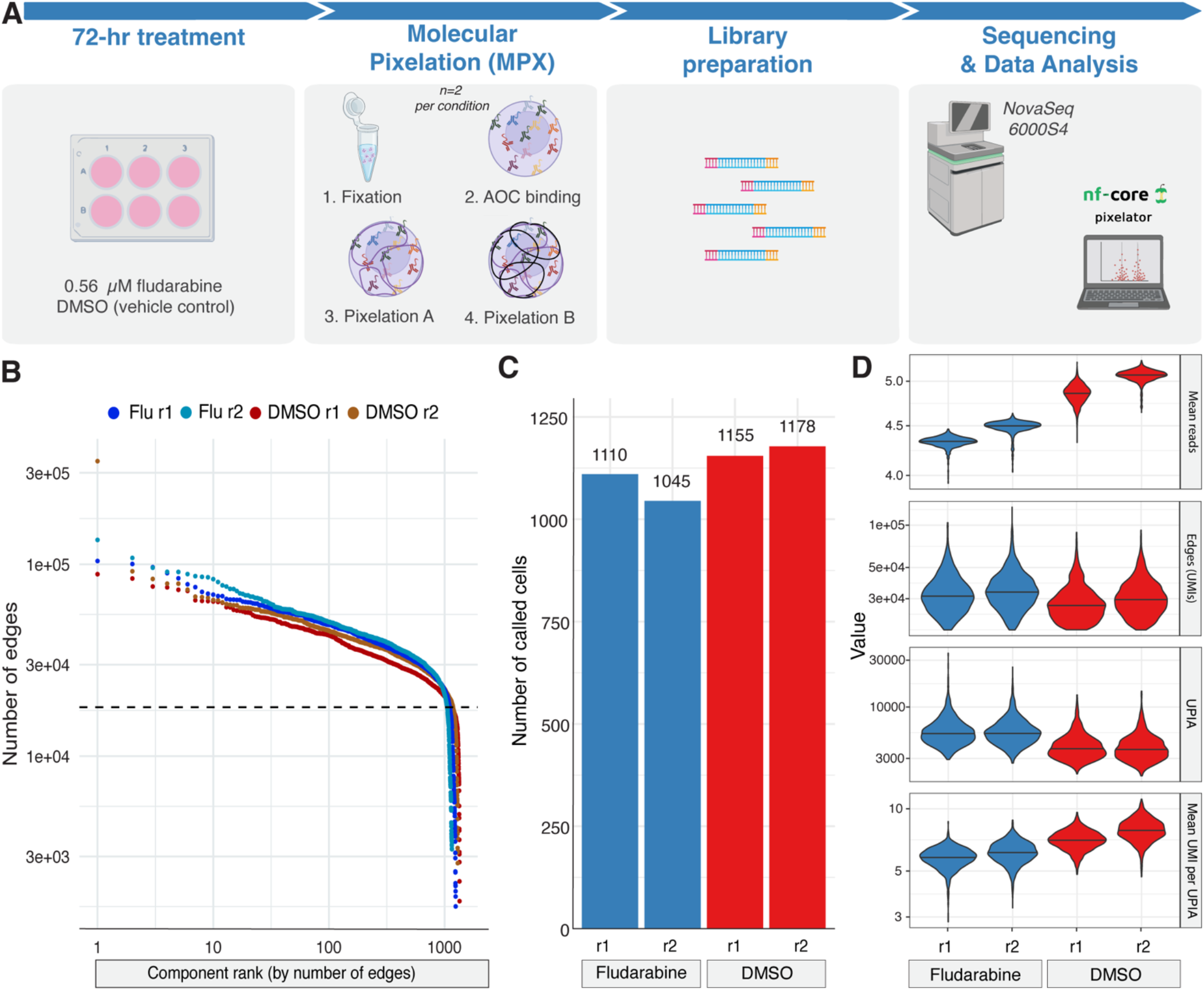
Study design and quality control metrics of fludarabine and DMSO treated Reh cells after Molecular Pixelation. **(A)** Graphical illustration of the study design. Reh cells were treated fludarabine or vehicle control for 72 hrs. Surviving cells were processed for single-cell spatial proteomics (Molecular Pixelation; MPX) in duplicate which includes a fixation step, an incubation with 80 antibody oligo conjugates (AOCs), two pixelation steps, a next generation sequencing (NGS) library preparation, sequencing, and data analysis on the nf-core/pixelator pipeline. **(B)** Edge-rank plot with number of edges (y-axis) and components (single-cells) ranked by number of edges (x-axis). A manual threshold (dashed black line) was set to filter out low-quality cells that deviate from the component size distribution and might not represent whole cells. **(C)** A bar chart showing the number of called cells per condition and replicate after a threshold was set on the edge-rank plot in (B). **(D)** Violin plots showing the number of mean reads, edges (also called Unique Molecular Identifier’s (UMIs)), DNA-pixels A, (UPIA) and mean UMI’s per UPIA in each condition and replicate. Figure 1A was created in BioRender (Globisch, M. (2024) BioRender.com/c77f038) and the MPX overview image was adapted from Karlsson *et al*., 2024^16^.

### Fludarabine induces changes in protein abundance and polarity

Next, we investigated differences in abundance and polarity. Using UMAP visualization, we noted differences between Reh cells after fludarabine treatment and DMSO (Figure 2A). Twenty-five proteins exhibited differences in abundance, with 23 increasing in expression and 2 decreasing in expression post-treatment (Figure 2B, Supplementary Table 7, average log2 Fold Change (log2FC) ≥ |0.1| and BH adj. p-value < 0.05). The proteins CD71 and CD44 decreased in abundance after fludarabine treatment, while HLA-DR, CD40, CD72, CD22, CD53, CD45, and CD82 increased in abundance (Figure 2C). The MPX polarity score indicates whether a protein is spatially clustered or dispersed on the cell surface^16^. We identified 24 proteins that increased in polarization post-treatment (Figure 2D-E, Supplementary Table 8, average polarity difference ≥ |0.1| and BH adj. p-value < 0.05). In total, 15 proteins (e.g. CD82, CD45, CD49D, CD47, HLA-DR, CD53, and CD40) changed in both abundance and polarity,10 proteins only changed in abundance (e.g. CD72), and 9 proteins only changed in polarity (e.g. CD38; Figure 2F).

**Figure 2.**
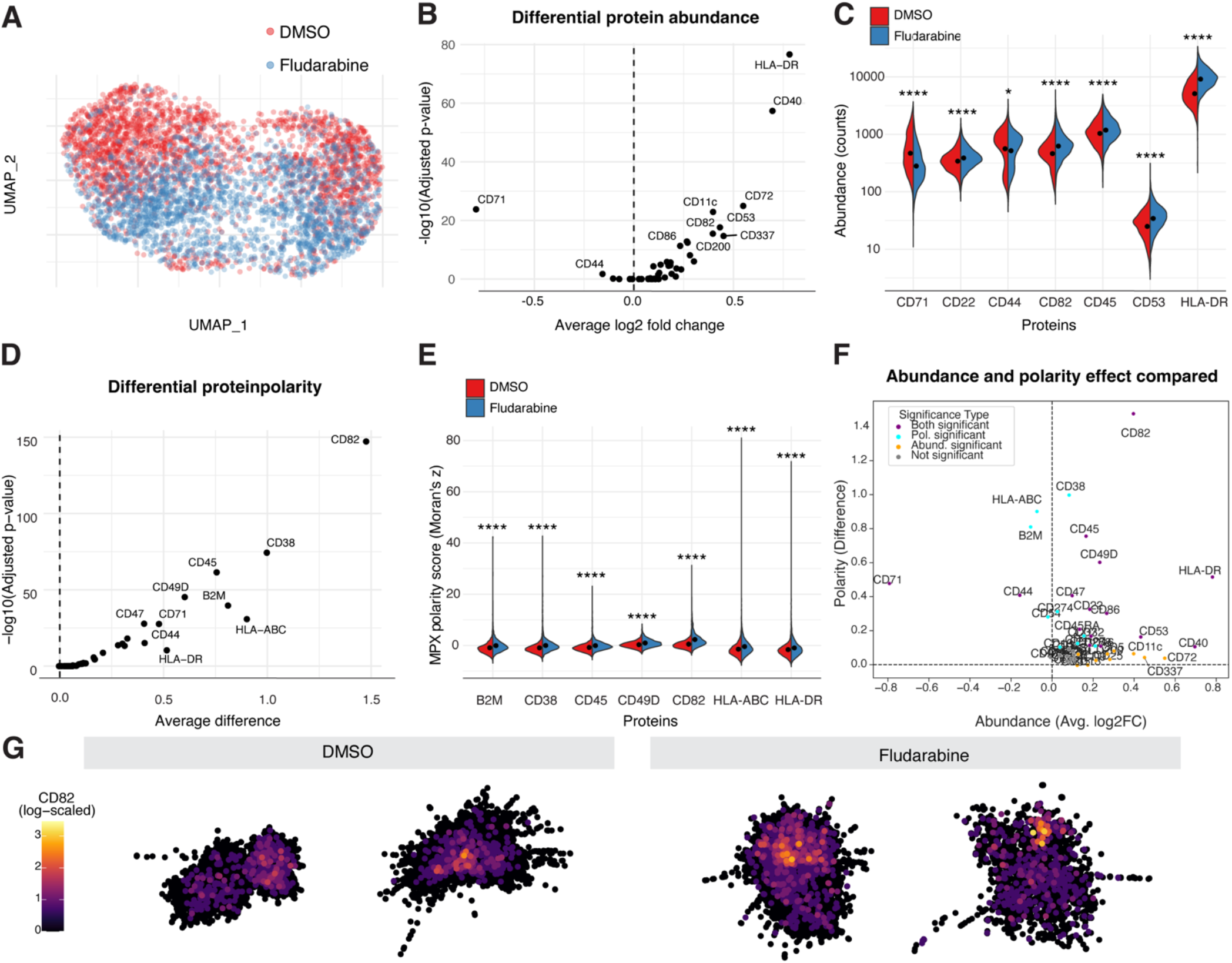
Immune surface proteins change in abundance and polarity in leukemia cells after fludarabine treatment. **(A)** A uniform manifold approximation and projection (UMAP) of Reh cells where counts from each cell (component) are clustered into two separate groups based on treatment; Fludarabine, blue and DMSO (control), red. **(B)** Volcano plot of differential protein abundance analysis (two-sided Wilcoxon Rank Sum test) of fludarabine treated cells compared with control cells. The y-axis depicts the Benjamin-Hochberg (BH) adjusted (adj.) p-value and the x-axis represents the average log2 fold change of each protein. The top statistically significant proteins are noted (average log2FC ≥ |0.1| and BH adj. p-value < 0.05). The dashed line divides the up-regulated (right) from the down-regulated (left) proteins. **(C)** Violin plots showing the abundance counts (y-axis) of selected differentially expressed proteins (x-axis) **(D)** Volcano plot of the differential protein polarity analysis (two-sided Wilcoxon Rank Sum test) of fludarabine treated cells compared to control cells (DMSO). The y-axis depicts the adj. p-value and the x-axis represents the average difference of each protein. The top statistically significant polarized proteins are noted (average polarity difference ≥ |0.1| and BH adj. p-value < 0.05) **(E)** Violin plots showing the MPX polarity score in Moran’s Z (y-axis) of selected differentially polarized proteins (x-axis) **(F)** Plot of abundance and polarity effect compared. Thresholds were set for abundance log2FC > |0.1|, polarity difference (> |0.1|), and adj. p-value < 0.05. **(G)** 2-dimensional MPX graphical representations of two selected cells per condition using Kamada-Kawai layout to illustrate the polarization of CD82. Each graph represents a single cell and each point on the graphs represent an individual DNA pixel A and is colored in proportion to the log10(counts+1) detected in the pixel.

Notably, CD82 and CD49D changed from a random spatial distribution on the cell surface in DMSO control cells (median polarity score of ∼0; Moran’s Z) to polarized in fludarabine treated cells (median polarity score > 0.5; Moran’s Z), suggesting that those two proteins become clustered after treatment. (Supplementary Figure 3; Supplementary Table 8). Two sampled cells from each condition were used to visualize the expression of CD82 in 2-dimensions (Figure 2G) and two additional sampled cells (1 for DMSO and 1 for fludarabine) were used to visualize the expression of CD82 in 3-dimensions (Supplementary Video 1 and 2, respectively).

### Fludarabine induces immunoprotein colocalization in leukemia cells

Protein colocalization refers to the spatial overlap of two or more proteins on cellular compartments, a characteristic that is necessary for signal transduction and cell migration^23^. The MPX colocalization score reflects the degree of spatial co-occurrence (positive score) or segregation (negative score) of two proteins^16^. We identified 138 protein pairs that changed in colocalization score upon treatment (Figure 3A-B, Supplementary Table 9; colocalization difference < 0.1 and BH adj. p-value < 0.05). CD82 colocalized more with CD49D and CD53, but segregated from numerous proteins including CD40, CD44, and CD45 in the fludarabine treated cells. Overall, CD82 stood out of the colocalization analysis (Figure 3C).

**Figure 3.**
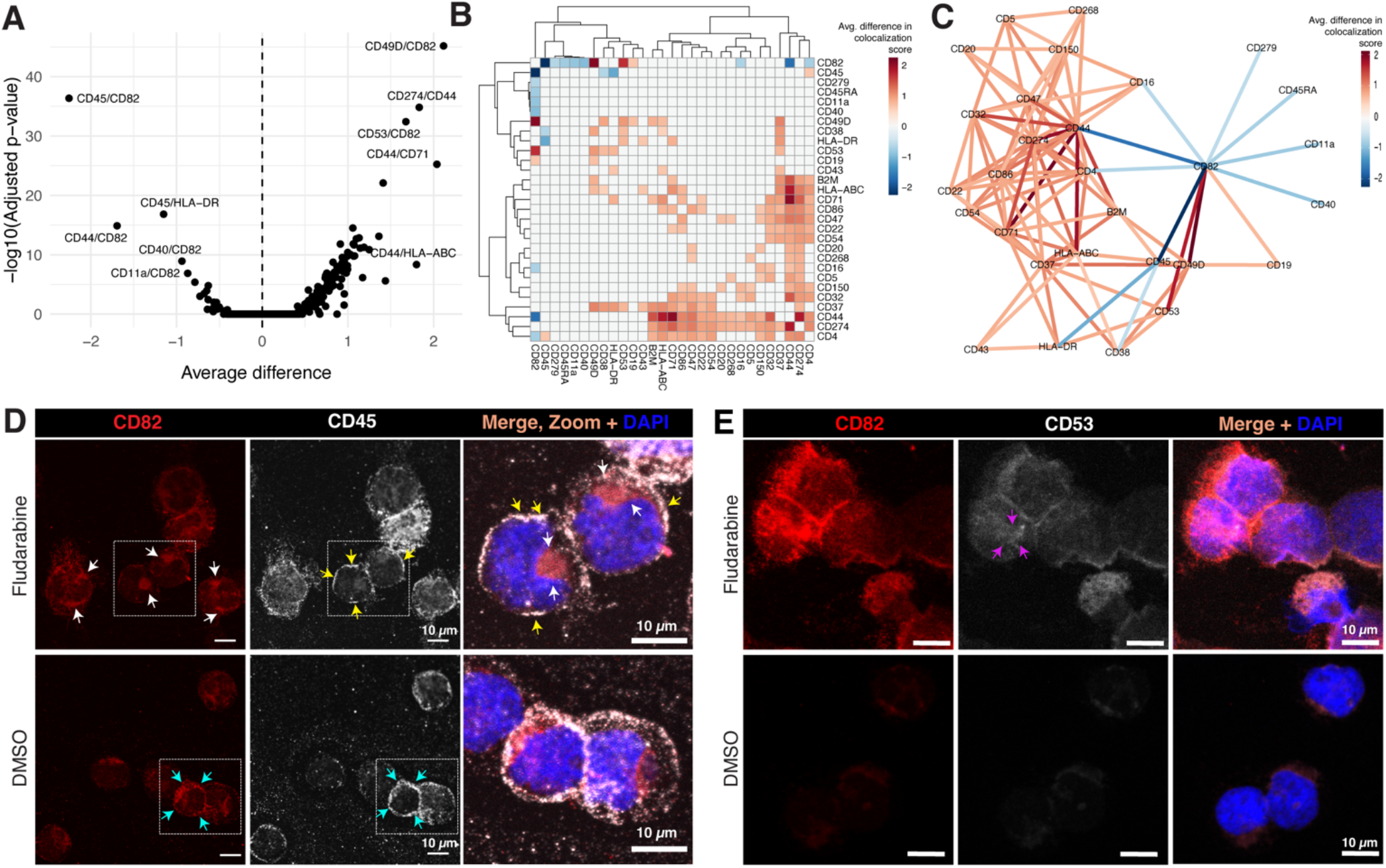
The spatial distribution of immune cell surface proteins changes in colocalization in Reh cells after fludarabine treatment. **(A)** Volcano plot of differential colocalization analysis of fludarabine treated cells compared to control cells (DMSO) The y-axis depicts the Benjamin-Hochberg (BH) adjusted (adj.) p-value and the x-axis represents the average difference between protein pairs. The top ten statistically significant pairs are noted (BH adj. p-value < 0.05; two-sided Wilcoxon Rank Sum test). **(B)** Heatmap with the average difference in colocalization score in protein pairs after fludarabine treatment. Red indicates higher colocalization and blue represents lower colocalization score. **(C)** Colocalization network illustrating the relationship between differentially colocalized proteins from A and B. Protein pair links are colored by the average difference in colocalization scores in Reh cells treated with fludarabine compared with control. **(D)** Immunocytochemistry confocal images of Reh cells treated with fludarabine or vehicle control (DMSO). Cells were stained for CD82 (red) and CD45 (white). DAPI (blue) was used to detect nuclei. The increased polarization of CD82 is noted with white arrows. **(E)** Immunocytochemistry confocal images of Reh cells treated with fludarabine or vehicle control (DMSO). Cells were stained for CD82 (red) and CD53 (white). DAPI (blue) was used to detect nuclei. The increased colocalization of CD82 and CD53 is noted with yellow arrows.

### ICC validates the change in abundance, polarization and colocalization of CD82, CD54, and CD45

Using an orthogonal ICC approach, we confirmed the increased abundance of CD82, CD45, and CD53 in fludarabine treated cells (Figure 3D-E). We also confirmed the increased polarization of CD82, CD45 and CD53 (white, yellow, and magenta arrows, respectively; Figure 3D-E) and the increased colocalization of CD82 with CD53 (Figure 3E, merged image; magenta). The few DMSO control treated cells that expressed CD82 and CD45 displayed the two proteins on the cell membrane (Figure 3D, cyan arrows) suggesting that the decreased colocalization of CD82 and CD45 observed in the fludarabine treated cells (MPX data) was due to the redistribution of CD82 and CD45 to different regions of the cell membrane (white and yellow arrows, respectively; Figure 3D, merged image). We also stained control and fludarabine treated leukemia cells with CD19, which did not change in abundance post treatment in the MPX data (Supplementary Table 7). These findings were further confirmed by ICC, which showed that the abundance of CD19 remained unaltered after fludarabine treatment, while CD82 and CD45 increased in the same post-treatment cells. In the MPX data, CD19 became polarized after treatment (Supplementary Table 8), however the increased polarization of CD19 in the fludarabine treated cells was not detectible in the ICC experiments (Supplementary Figure 4).

### scRNA-seq counts and cell surface protein expression

We previously identified transcriptomic changes in Reh cells post fludarabine treatment by scRNA-seq^15^. Herein, we used these data to compare the transcript levels to cell surface protein abundance measurements from the MPX assay. Using the transformed pseudo-bulk counts, we compared the difference in expression between fludarabine and DMSO control cells (MAST test; Supplementary Table 10). We selected the genes coding for the 76 cell surface proteins (Supplementary Table 2), identifying 49 genes expressed in the scRNA-seq Reh dataset (Supplementary Figure 5A-B). Of these genes, 18 were differentially expressed (16 up- and 2 down-regulated; log2FC > 0.50; adj. p-value < 0.01; MAST test) (Figure 4A and Supplementary Figure 5A-B). We subsequently compared the 18 DEGs (scRNA-seq) with the 25 differentially expressed proteins (MPX) and found consensus in significance and directionality for nine genes/proteins including CD40, *PTPRC*/CD45, CD53, and CD82 (Figure 4B and Supplementary Table 11). Overall, there was a positive correlation between the changes in transcript levels and cell surface protein expression in fludarabine treated and DMSO control cells (*ρ*= 0.382; Figure 4C).

**Figure 4.**
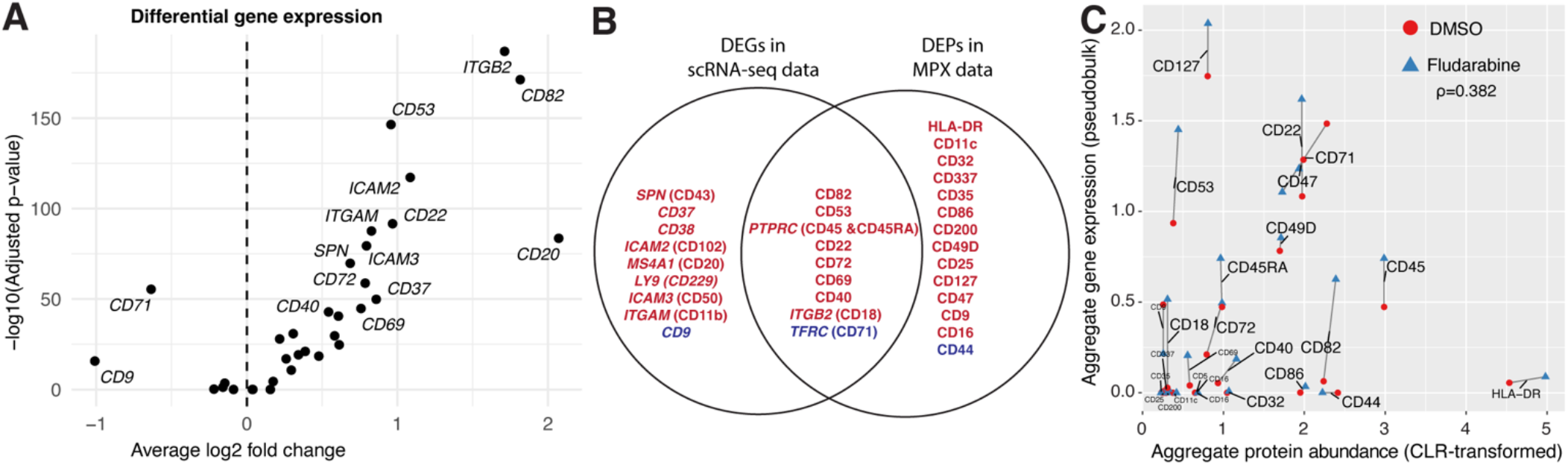
scRNA-seq counts and cell surface protein expression. **(A)** Volcano plot illustrating the differentially expressed genes identified in fludarabine treated Reh cells compared to control cells (DMSO). The y-axis depicts the -log10 adjusted (adj.) p-value and the x-axis represents the average log2 fold change. The top statistically significant pairs are noted (log2FC > 0.50; adj. p-value < 0.01; MAST test). **(B)** Ven Diagram illustrating the overlap between gene and protein expression patterns of cell surface markers identified between fludarabine and control treated cells. Red is upregulated and blue is downregulated. **(C)** Correlation plot showing aggregate protein abundance as centered log ratio (CLR)-transformed (x-axis, MPX data) and aggregate gene expression data as pseudobulk (y-axis, scRNA-seq data) of the 25 differentially expressed proteins (x-axis, adj p-value less < 0.05) identified by MPX between fludarabine and DMSO control cells. A Pearson analysis was used to determine the correlation of the fludarabine/DMSO log2FC in both scRNAseq and MPX. The correlation coefficient (*ρ*) is noted. Gene expression data from Gazelius *et al*., 2024 (accession number GSE229617) were used in A, B, and C and in B and C, scRNAseq data were combined with MPX data.

## Discussion

Spatial biology is an emerging field that provides new insights into the spatial organization and functional dynamics of cells and tissues^24^. In this study, we leveraged MPX, a state-of-the-art single-cell spatial proteomics assay^16^ to examine changes in the cell surface proteome of leukemia cells following exposure to fludarabine, a chemotherapeutic agent commonly used in the treatment of advanced hematologic malignancies. By profiling the spatial distribution of 76 targeted surface proteins on individual cells, we investigated key aspects of cell surface dynamics, including protein abundance and polarity as well as colocalization of protein pairs to better understand the cellular response to fludarabine.

Key findings in our study include the increased abundance and polarization of CD82 and CD53 in surviving fludarabine-treated leukemia cells. CD82 and CD53 are members of the tetraspanin family and have established roles in receptor-mediated signaling, immune-regulation, cell adhesion, migration, growth and differentiation^25^. Notably, CD82 and CD53 have been implicated in immune cell modulation, hematopoietic stem cell (HSC) stress response and cancer suppression^26-28^. In acute myeloid leukemia (AML), CD82 and CD53 transcript and protein levels are elevated following daunorubicin chemotherapy in cell lines and patient-derived samples^29^. Interestingly, in the same study CD82 overexpression was linked to reduced apoptosis in response to chemotherapy, while CD82 knockdown restored chemosensitivity. In the Human Protein Atlas, CD82 expression is highly correlated with p53 expression, which mirrors our previous findings with scRNAseq^15^. The increased abundance and polarization of CD82 and CD53 observed in this study further suggests that these two tetraspanins have important roles in cellular stress response, survival mechanisms and adaptive responses to chemotherapy. Our findings, together with previous reports^30,31^, suggest that targeting CD82, CD53 or downstream signaling proteins could present a novel therapeutic strategy to mitigate chemoresistance in leukemia.

Therapy-induced cellular senescence is a state where cancer cells stop proliferating and resist growth stimuli following chemotherapy exposure^32^. While senescence halts proliferation, it may paradoxically promote cancer cell survival^33^. In our study, we observed that Reh cells treated with fludarabine were larger compared to control cells, possibly indicating cellular senescence^34^. Senescent cells often undergo changes that enhance their recognition by the immune system, potentially marking them for clearance. This observation, if also relevant in vivo, may partly explain some of the clinical benefits observed when fludarabine is used in preparation for CAR T-cell therapy^10^. Our results suggest that, beyond its well-documented lymphodepleting effects^35^, fludarabine may also induce immunomodulatory changes on the surface of leukemia cells, potentially priming them for more efficient immune-mediated destruction by CAR-T cells^36^. A key finding was the increased abundance and polarization of HLA-DR (MHC class II) and CD40 following fludarabine treatment. Both HLA-DR and CD40 play essential roles in activation of antigen-presenting cells^37^ and are expressed in primary ALL cells^38,39^. CD40 is of particular interest due to its potential as a therapeutic target in cancer immunotherapy^40,41^. Notably, CAR-T cells expressing CD40 ligand have demonstrated superior antitumor efficacy and enhance the recruitment of immune effectors in leukemia and lymphoma mouse models^42^.

The application of spatial proteomics^16^ in our study revealed novel insights into the adaptive changes in the cell surface proteome in response to fludarabine treatment. By leveraging the MPX assay, we were able to pinpoint alterations in the spatial organization of surface proteins, offering new insight on how leukemia cells respond to fludarabine. However, our study has several limitations. We analyzed only one leukemia cell line and focused on a subset of 76 surface proteins, which may not fully reflect the diversity and complexity of the leukemia cell surface proteome, particularly across the numerous distinctive molecular subtypes of ALL^43^. Future studies should include additional leukemia cell lines and, crucially, patient-derived samples to validate and extend these findings. Moreover, our analysis was restricted to a single timepoint (72 hrs post-treatment). Tracking the spatiotemporal dynamics of the cell surface proteome at multiple timepoints could provide even greater insights into how leukemia cells adapt during the course of fludarabine treatment, potentially uncovering further therapeutic opportunities.

In conclusion, our study provides the first in-depth characterization of the spatial organization of the surface proteome of leukemic cells. We identified changes in the expression and spatial organization of numerous cell surface proteins, with CD40, CD53 and CD82 emerging as key proteins in the cellular response to fludarabine. These findings deepen our understanding of leukemia cell biology in the context of fludarabine treatment and lay the groundwork for future investigations into the mechanisms driving chemotherapy resistance.

## Supporting information

Globisch et al 2024 Supplementary Material

Globisch et al 2024 Supplementary Tables

Globisch et al 2024 Supplementary Video 1

Globisch et al 2024 Supplementary Video 2

## Data availability

The scRNA-seq data of fludarabine and DMSO treated Reh cells are available at the Gene Expression Omnibus (GEO) under accession number GSE229617^15^. The raw sequencing data produced from single-cell spatial proteomics are available at the Sequence Read Archive under accession number PRJNA1167477.

## Author contributions

MAG: Conceptualization, methodology, validation, formal analysis, investigation, writing-original draft, visualization, project administration.

HG: Conceptualization, investigation, writing - review & editing

APE: Conceptualization, resources, writing - review & editing

AL: Software, formal analysis, data curation, visualization, writing - review & editing

OK: Formal analysis, visualization, writing - review & editing

MLW: Formal analysis, writing - review & editing

MÅ: Investigation

AH: Resources

AR: Resources, writing - review & editing

CA: Investigation, methodology, writing - review & editing

JN: Conceptualization, writing- original draft, supervision, project administration, funding acquisition

## Acknowledgements

This work was supported by grants from the Swedish Research Council (2019-01976 to JN), the Swedish Cancer Society (CAN2022-2395 to JN), the Swedish Childhood Cancer Foundation (PR2022-0082 and HFT2023-0011 to JN, TJ2020-0039 to AH), and the Swedish Society for Medical Research (PG-23-0420-H-01to MAG). We thank Pixelgen technologies for providing us with the opportunity to beta test their single-cell spatial proteomics assay, in particular Louise Leijonancker and Stefan Petkov for their assistance in training us to run the assay and bioinformatics analyses. We are grateful to Nina Rolf and Chinten James Lim for their valuable perspectives on the study. Assistance with sequencing was provided by the SciLifeLab National Genomics Infrastructure at Uppsala University, funded by SciLifeLab, the Knut and Alice Wallenberg Foundation, and the Swedish Research Council through grant agreement no. 2019-0222. We also thank the BioVis Core Facility at Uppsala University for technical assistance with the fluorescent microscopes. The computational analyses were supported by resources provided by the National Academic Infrastructure for Supercomputing in Sweden (NAISS), partially funded by the Swedish Research Council through grant agreement no. 2022-06725.

