## Supplementary material for "Mapping the Spatial Proteome of Leukemia Cells Undergoing Fludarabine Treatment": Globisch et al 2024 Supplementary Material

### Supplementary figures

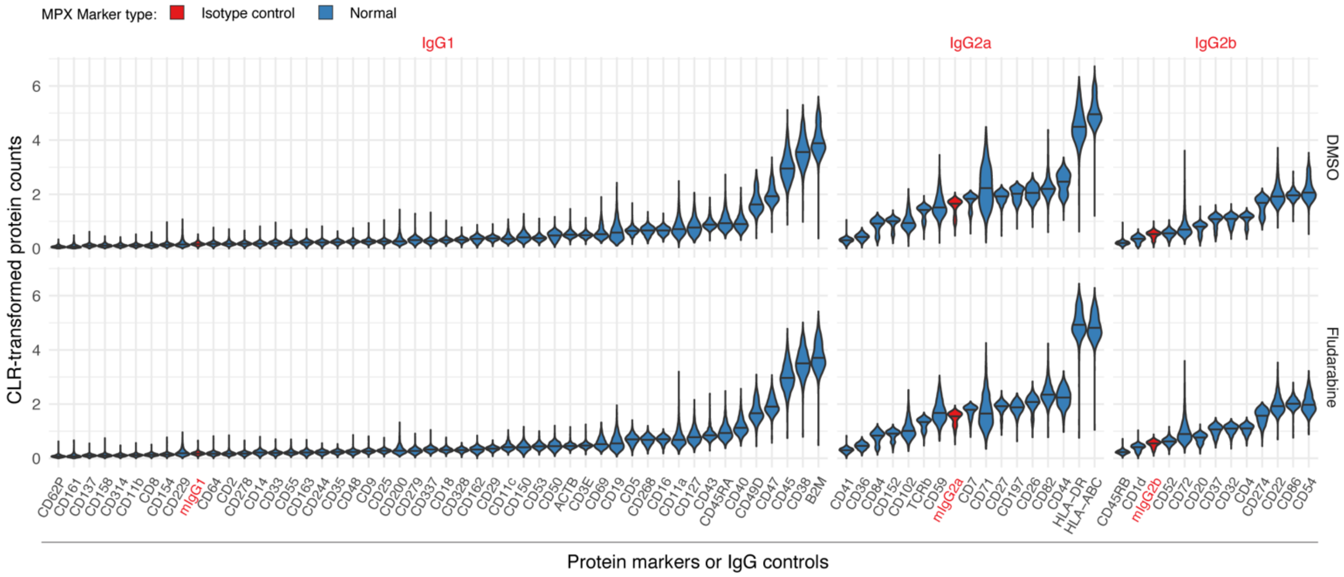


**Supplementary Figure 1. Overview of the most variable proteins and isotype controls.** Violin plots showing the centered log ratio (CLR)-transformed counts of the most variable proteins identified in Reh cells and the internal controls which are marked in red (IgG1, IgG2a, and IgG2b).

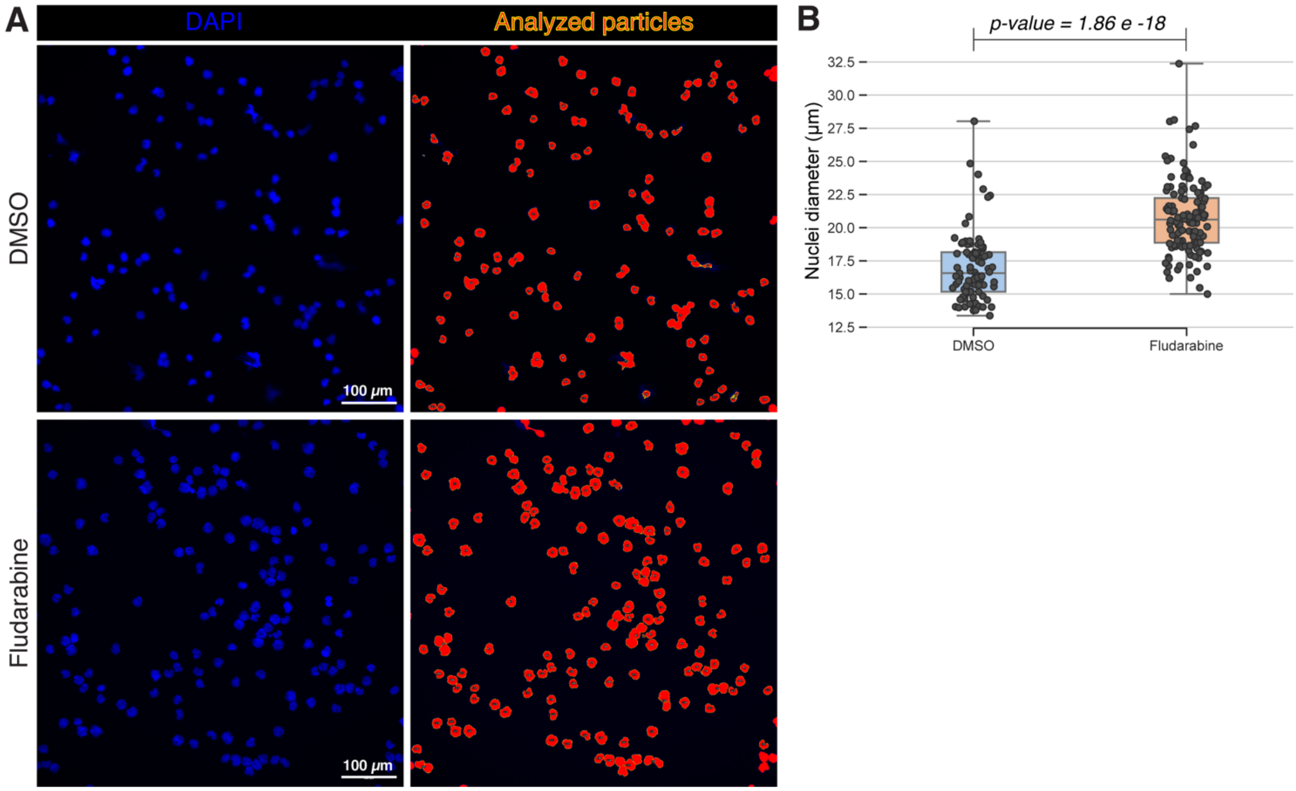


**Supplementary Figure 2. Nuclear measurements of leukemia cells after fludarabine and control treatments. (A)** Immunofluorescent images of Reh cells treated with DMSO (control) or fludarabine and then stained with DAPI (blue) to identify nuclei. A threshold was set on the DAPI signal (red) and the *analyze particles* function was used on Image J to select all nuclei per condition (red, yellow outline). **(B)** The diameters of selected nuclei were calculated. A two-sided t-test was used to assess the difference between DMSO and fludarabine treated cells.

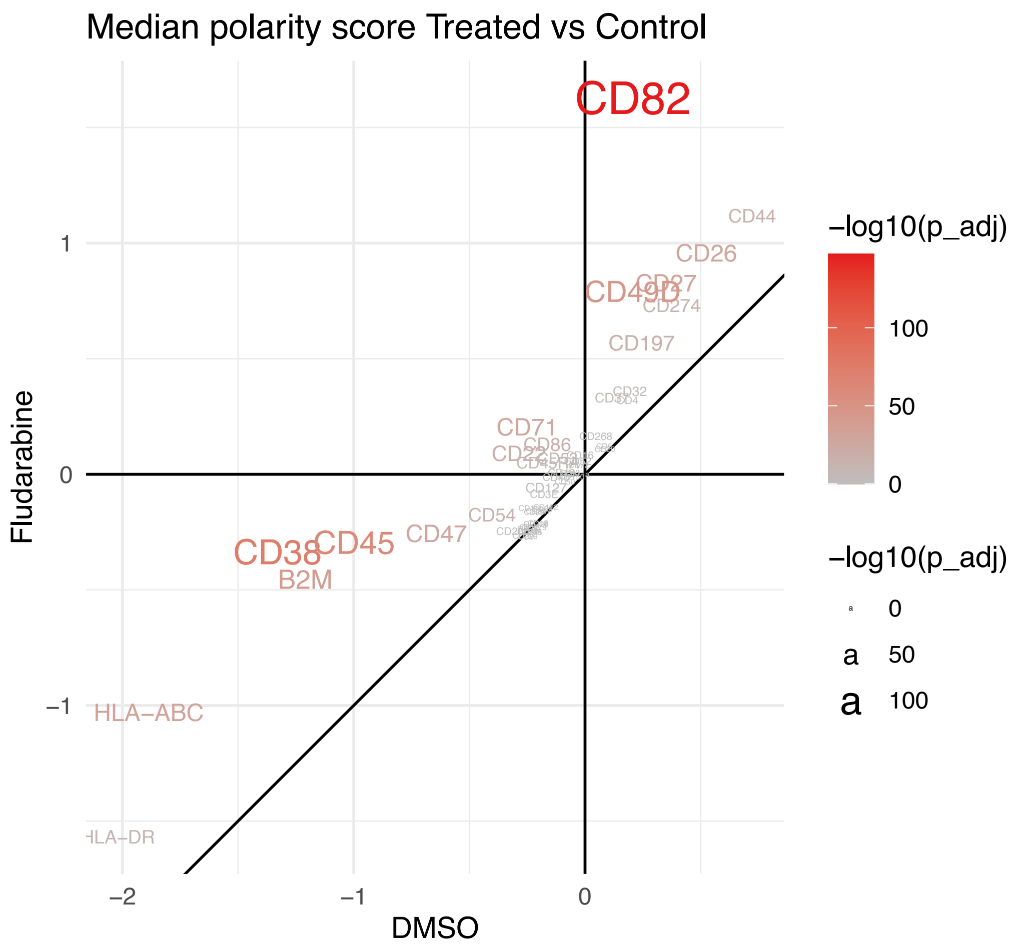


**Supplementary Figure 3. Polarized proteins in leukemia cells after fludarabine treatment.** Scatter plot illustrating the median polarity scores (Moran's Z) of polarized proteins in DMSO (x-axis) and fludarabine (y-axis) treated leukemia cells. The color and size of the protein marker represents the -log10 of the adjusted p-value (p_adj).

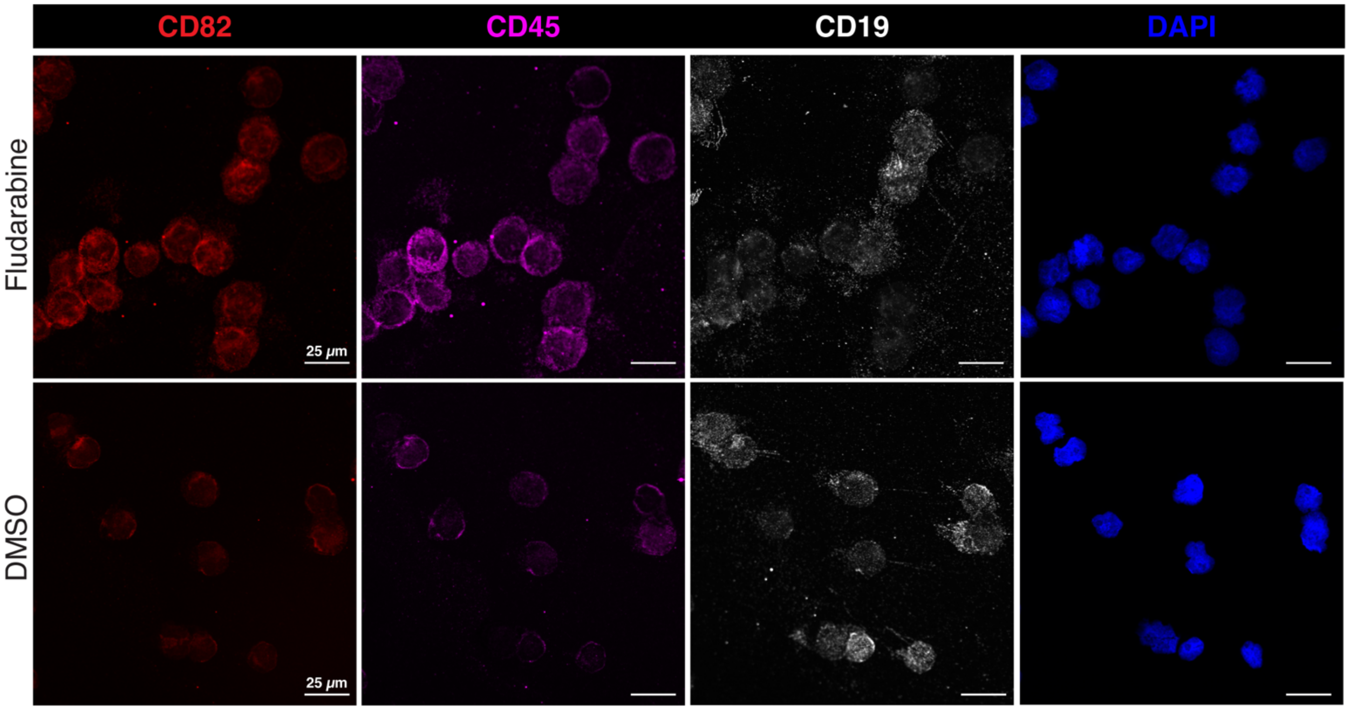


**Supplementary Figure 4. Immunocytochemistry of cell surface proteins in fludarabine and control treated leukemia cells.** Confocal images of Reh cells treated with fludarabine or vehicle control (DMSO). Cells were stained for CD82 (red) and CD45 (magenta) and DAPI (blue) was used to detect nuclei.

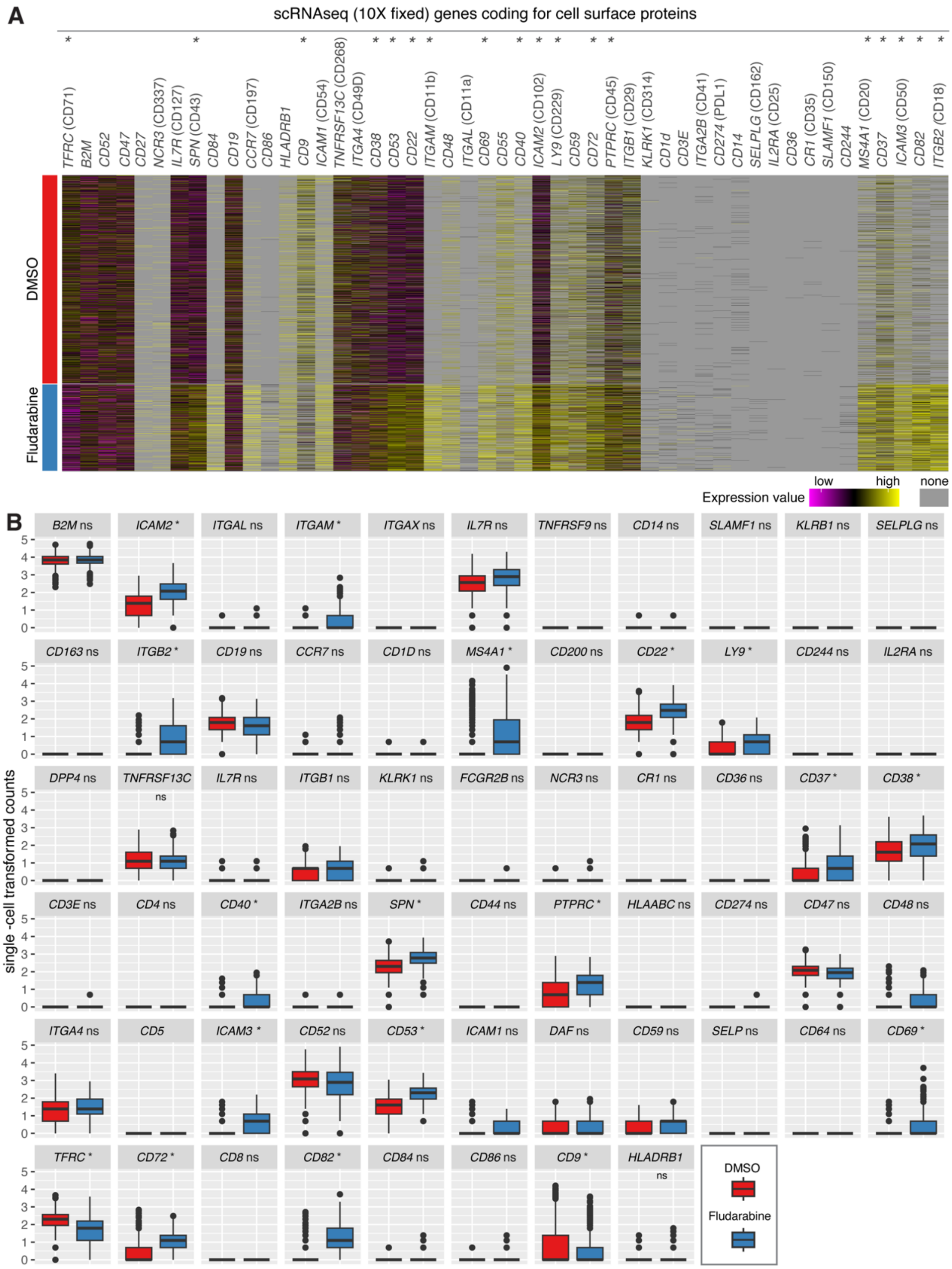


**Supplementary Figure 5. Transcript levels of genes coding for cell surface proteins. (A)** Heatmap illustrating the single-cell transformed and normalized counts in DMSO and fludarabine treated Reh cells. Genes are listed on the top of the heatmap with surface protein in parentheses if the protein name is different than the gene name. **(B)** Boxplots showing the single-cell transformed and normalized counts of genes that code for cell surface proteins. In the heatmap and boxplots differentially expressed genes are marked with an asterisk (log2FC > 0.50; adj. p-value < 0.01; MAST test). In the boxplots, only the gene names are noted and non-differentially expressed genes are marked with ns for not significant.

**Supplementary video 1. Three-dimensional representation of CD82 expression in a DMSO control leukemia cell.** A DMSO control leukemia cell expressing CD82 was used to visualize the spatial localization of CD82 in DMSO control conditions. Each point represents an individual DNA pixel A.

**Supplementary video 2. Three-dimensional representation of CD82 expression in a fludarabine treated leukemia cell.** A fludarabine treated leukemia cell expressing CD82 on the cell surface was used to visualize the spatial localization of CD82 in fludarabine conditions. Each point represents an individual DNA pixel A.
